# A distinctive ligand recognition mechanism by the human vasoactive intestinal polypeptide receptor 2

**DOI:** 10.1101/2021.09.16.460712

**Authors:** Yingna Xu, Wenbo Feng, Qingtong Zhou, Anyi Liang, Jie Li, Antao Dai, Fenghui Zhao, Lihua Zhao, Tian Xia, Yi Jiang, H. Eric Xu, Dehua Yang, Ming-Wei Wang

## Abstract

Activated by physiologically important peptide hormones, class B1 G protein-coupled receptors (GPCRs) modulate key physiological functions and serve as valuable drug targets for many diseases. Among them, vasoactive intestinal polypeptide receptor 2 (VIP2R) is the last member whose full-length 3-dimensional structure has yet to be determined. VIP2R, expressed in the central and peripheral nervous systems and involved in a number of pathophysiological conditions, is implicated in pulmonary arterial hypertension, autoimmune and psychiatric disorders. Here, we report the cryo-electron microscopy structure of the human VIP2R bound to its endogenous ligand PACAP27 and the stimulatory G protein. Different from all reported peptide-bound class B1 GPCR structures, the N-terminal α-helix of VIP2R adopts a unique conformation that deeply inserts into a cleft between PACAP27 and the extracellular loop 1, thereby stabilizing the peptide-receptor interface. Its truncation or extension significantly decreased VIP2R-mediated cAMP accumulation. Our results provide additional information on peptide recognition and receptor activation among class B1 GPCRs and may facilitate the design of better therapeutics.

## Introduction

Vasoactive intestinal peptide (VIP) and pituitary adenylate cyclase-activating polypeptide (PACAP) are two important neuropeptides that exert a variety of physiological actions through three class B1 G protein-coupled receptors (GPCR), namely PACAP type 1 receptor (PAC1R), VIP receptors 1 (VIP1R, or VPAC_1_) and 2 (VIP2R, or VPAC_2_)^1^. They share about 50% sequence similarities but mediate different functions such as neural development, calcium homeostasis, glucose metabolism, circadian rhythm, thermoregulation, inflammation, feeding behavior, pain, stress and related endocrine responses^2-6^. Interestingly, PACAP (PACAP38 and PACAP27, a C-terminally truncated variant of PACAP38) and VIP have comparable affinity at VIP1R and VIP2R, but PACAP is 400 to 1000-fold more potent than VIP at the PAC1R (Fig. 1a, b). Extensively expressed in the central and peripheral nervous systems^7, 8^, VIP2R is involved in a number of pathophysiological conditions, showing a great potential as a therapeutic target for pulmonary arterial hypertension, chronic obstructive pulmonary disease (COPD), cancer, asthma, autoimmune and psychiatric disorders^9-12^.

**Figure 1.**
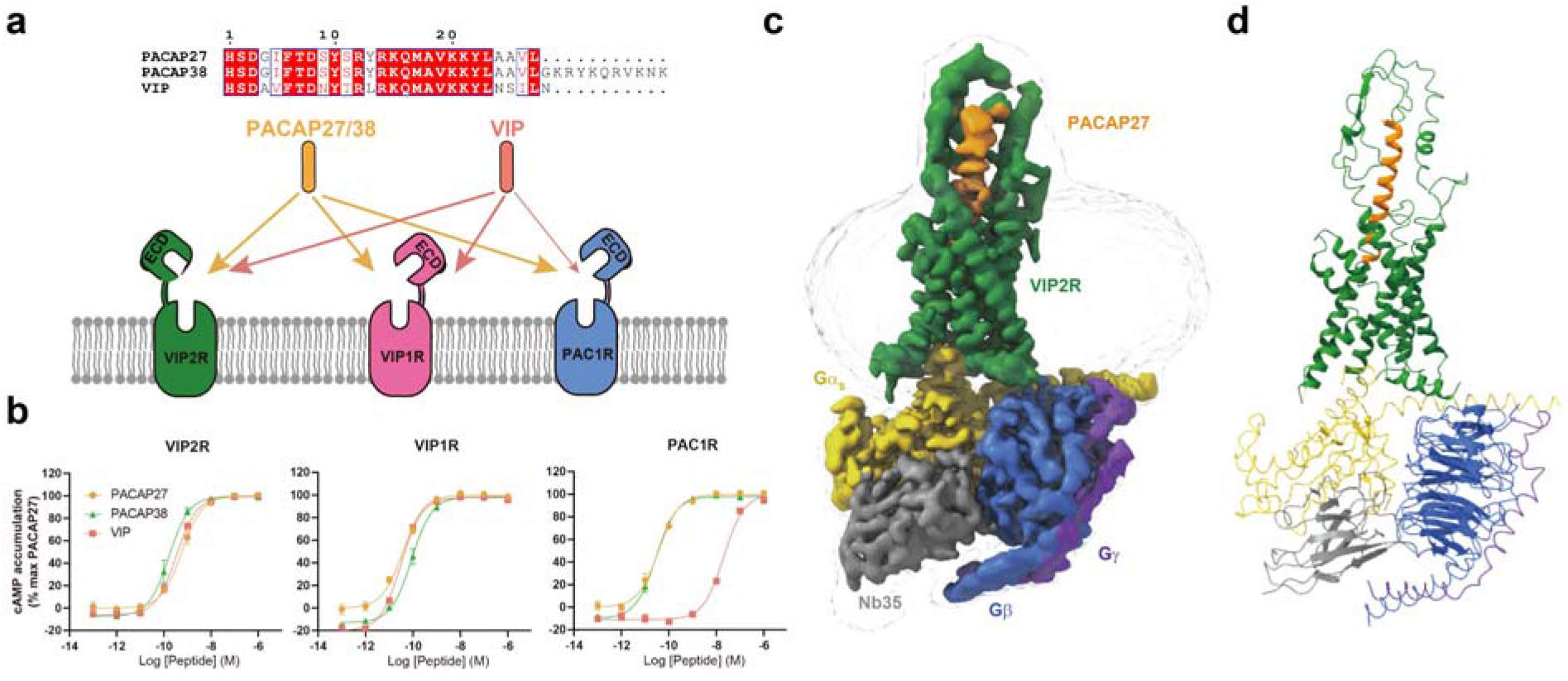
Cryo-EM structure of the PACAP27–VIP2R–G_s_ complex. **a**, Binding specificity of PACAP and VIP receptor subfamily to the related peptide hormones. Sequence alignment of peptides are shown on the top panel. **b**, Receptor signaling profiles of endogenous agonists PACAP27, PACAP38 and VIP. Data shown are means ± S.E.M. of three independent experiments performed in quadruplicate. **c**, Cut-through view of the cryo-EM density map illustrating the PACAP27–VIP2R–G_s_ complex and the disc-shaped micelle. **d**, Model of the complex as a cartoon, with PACAP27 as helix in orange. The receptor is shown in green, Gα_s_ in yellow, Gβ subunit in royal blue, Gγ subunit in violet, and Nb35 in gray.

A comprehensive molecular understanding of VIP/PACAP recognition and receptor activation is important to the design of better drug candidates. Further to the recently reported cryogenic electron microscopy (cryo-EM) structures of VIP1R and PAC1R^13-16^, we determined a single-particle cryo-EM structure of the human VIP2R in complex with PACAP27 and the stimulatory G protein (G_s_), at a global resolution of 3.4 Å. Combined with molecular dynamics (MD) simulation and functional studies, we obtained some valuable insights into a distinctive molecular mechanism governing ligand recognition and VIP2R activation.

## Results

### Structure determination

To prepare a high quality human VIP2R–G_s_ complex, we added a double tag of maltose binding protein at the C terminus, replaced the native signal peptide at the N terminus with the prolactin precursor sequence (Supplementary Fig. 1a–b), and in employed the NanoBiT tethering strategy^13, 17-20^. The resultant VIP2R construct retained the full-length receptor sequence (residues 24-438 excluding the signal peptide sequence), different from that of VIP1R^13^ and PAC1R^14, 15^ where either C-terminal truncation, mutations or their combination were made. Large-scale purification was followed and the PACAP27–VIP2R–G_s_ complexes were collected by size-exclusion chromatography (SEC) for cryo-EM study. (Supplementary Fig. 1c–e). After sorting by constitutive 2D and 3D classifications, 3D consensus density maps were reconstructed with a global resolution of 3.4 Å (Fig. 1c, Supplementary Fig. 2 and Supplementary Table 1). The cryo-EM maps allowed us to build an unambiguous model for most regions of the complex except for the flexible α-helical domain (AHD) of Gα_s_ and the VIP2R residues from P313 to S321 in the intracellular loop 3 (ICL3) (Supplementary Fig. 3). The extracellular domain (ECD) had a lower resolution due to high conformational flexibility widely observed among class B1 GPCRs^21-23^, and therefore was modelled on the rigid-body fitted crystal structure of VIP2R ECD (PDB code: 2×57).

### Overall structure

The PACAP27–VIP2R–G_s_ complex adopts a typical architecture of the activated class B1 GPCR conformations, characterized by a single straight helix of PACAP27 that interacts with both ECD and the transmembrane domain (TMD), a sharp kink in the middle of the TM helix 6 (TM6) thereby opening the cytoplasmic face, and the insertion of the C-terminal α5 helix of the Gα_s_ into the receptor core (Fig. 1c). Its overall structure is similar to other class B1 GPCRs–G_s_ complexes such as GLP-1–GLP-1R–G_s_ (PDB code: 6×18)^24^, GIP–GIPR–G_s_ (PDB code: 7DTY)^25^, TIP39–PTH2R–G_s_ (PDB code: 7F16)^26^ and UCN1–CRF1R–G_s_ (PDB code: 6PB0)^27^ with root mean square deviation (RMSD) values of 1.40, 0.99, 1.39, and 0.96 Å for the whole complex, respectively. Meanwhile, the structure of PACAP27-bound VIP2R displays a high degree of similarity compared to both PACAP27-bound VIP1R (PDB code: 6VN7)^13^ and PAC1R bound by PACAP38 (PDB code: 6P9Y)^16^ and maxadilan (PDB code: 6M1H)^14^, with Cα RMSD of 0.70, 0.96 and 1.04 Å, respectively.

As shown in Figure 2, the N termini of the bound PACAP27 and PACAP38 overlapped well and penetrated into the TMD core by an almost identical angle and orientation, exhibiting a shared ligand recognition pattern (Supplementary Table 2). Notable differences in both position and orientation at the peptide C-terminal halves were observed via the surrounding ECD, ECL1 and the extracellular tip of TM1 conformations that are unique to VIP2R. Specifically, the VIP2R-bound PACAP27 was rotated by 4.6° compared to that in complex with VIP1R; such a movement shifted its C terminus toward the TMD core by 4.3 Å (measured by the Cα of L27^P^, P indicates that the residue belongs to the peptide). By choosing a more relaxed ECL1 conformation rather than the ordered two-turn α-helix found in the ECL1 of PAC1R, VIP2R reduced the contacts between ECL1 and peptide evidenced by a decrease in the buried surface area from 282 Å^2^ (PAC1R) to 77 Å^2^ (VIP2R). Consequently, the C terminus of PACAP27 bound by VIP2R moved toward ECL1 by 2.6 Å in comparison with that of PACAP38 bound by PAC1R (measured by the Cα of L27^P^). Collectively, these common and unique structural features highlight the complexity of peptide recognition among VIP2R, VIP1R and PAC1R.

**Figure 2.**
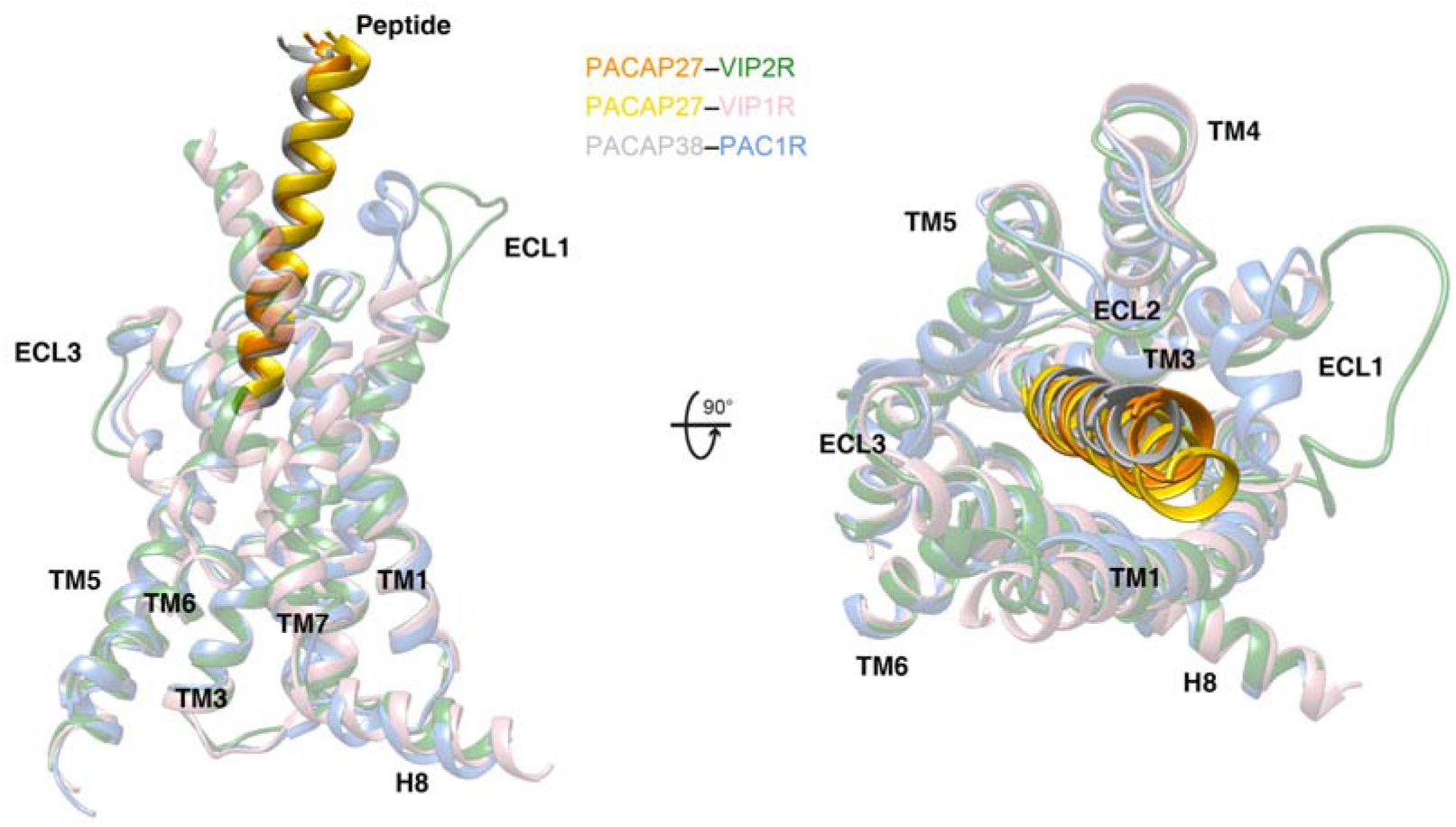
Structural comparison of active VIP2R, VIP1R and PAC1R. Superimposition of PACAP27–VIP2R, PACAP27–VIP1R (PDB code: 6VN7)^13^ and PACAP38–PAC1R (PDB code: 6M1I)^14^reveals a high structural similarity. Receptor ECD and G protein are omitted for clarity.

### Ligand recognition

The active VIP2R structure shows that PACAP27 is stably anchored through two interaction networks (Fig. 3a): the first is comprised of the peptide N-terminal half (residues 1 to 13) and the residues in the lower half of the ligand-binding pocket (Fig. 3b), while the second connects the peptide C-terminal half with the ECD (especially the N-terminal α-helix), ECL1 and the stalk region (Fig. 3c).

**Figure 3.**
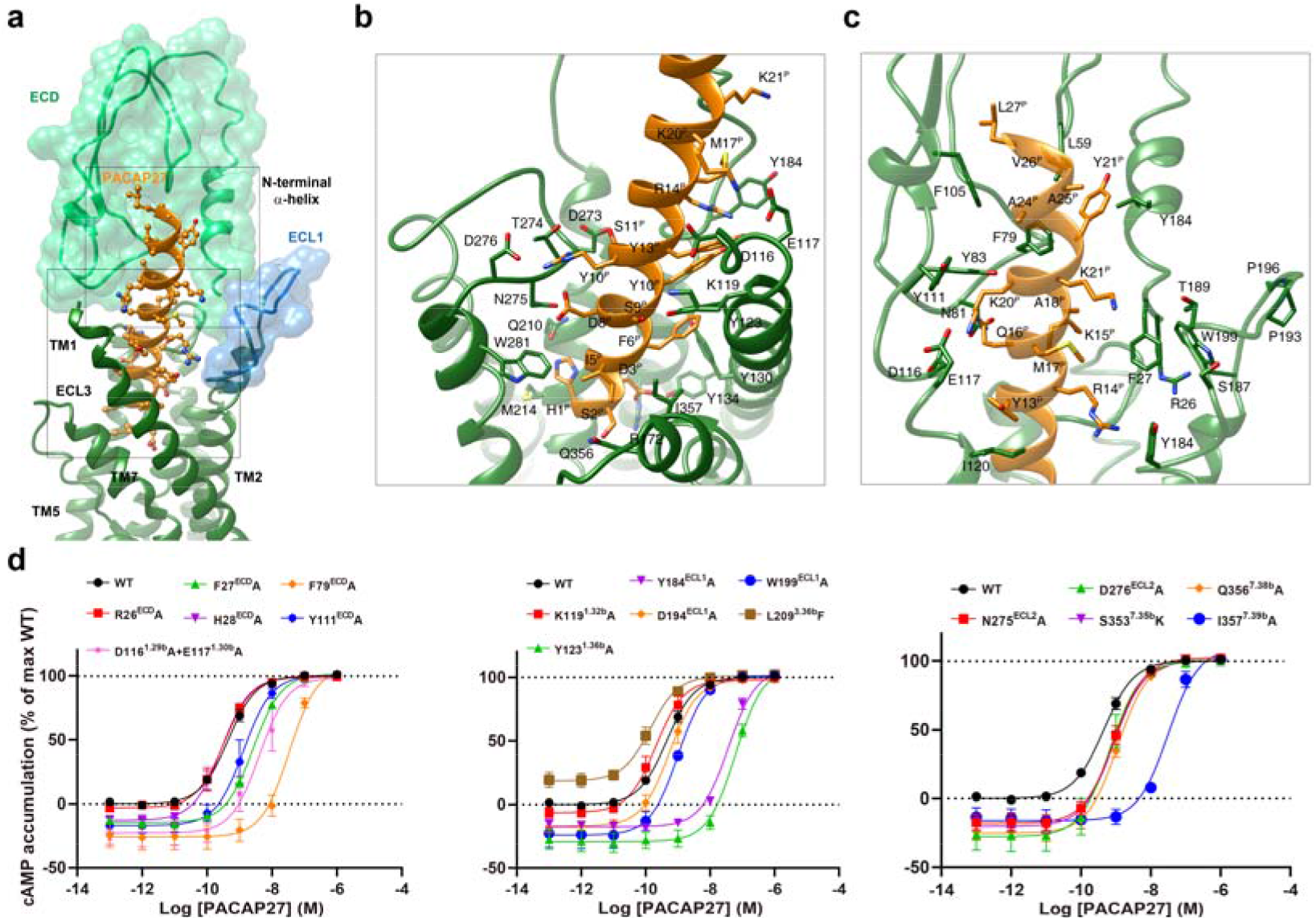
Molecular recognition of PACAP27 by VIP2R. **a**, The binding mode of PACAP27 (orange) with VIP2R (green), showing that the N-terminal half of PACAP27 penetrates into a pocket formed by TMs 1-3, TMs 5-7 and ECLs 1-3, whereas the C-terminal half is recognized by the ECD, ECL1 and TM1. The ECD and ECL1 are shown as surface. **b**, Close-up view of the interactions between PACAP27 and the TMD pocket of VIP2R. **c**, Close-up view of the interactions between the C-terminal half of PACAP27 and the peptide-binding pocket of VIP2R. Key residues are shown as sticks. **d**, Signaling profiles of VIP2R mutants. cAMP accumulation in wild-type (WT) and single-point mutated VIP2R expressing in CHO-K1 cells. Signals were normalized to the maximum response of the WT and dose–response curves were analyzed using a three-parameter logistic equation. All data were generated and graphed as means ± S.E.M. of at least three independent experiments, conducted in quadruplicate.

In the first network, H1^P^, D3^P^, I5^P^, F6^P^, D8^P^, S9^P^, Y10^P^ and S11^P^ contribute common interactions with the conserved residues among VIP2R, VIP1R and PAC1R, while S2^P^, G4^P^, R12^P^ and Y13^P^ make receptor-specific interactions. For common interactions, H1^P^ is oriented toward TM3 with the formation of a hydrogen bond with Q210^3.37b^ (class B GPCR numbering in superscript), similar interaction is also observed in VIP1R and PAC1R. D3^P^ is highly conserved in class B1 GPCR peptide hormones, which simultaneously forms one salt bridge with R172^2.60b^ and one hydrogen bond with Y134^1.47b^ in the cases of VIP2R, VIP1R and PAC1R. Polar interactions also occurred between D8^P^ and N275^ECL2^, S9^P^ and Y123^1.36b^, as well as S11^P^ and D273^ECL2^. I5^P^, F6^P^ and Y10^P^ contribute massive nonpolar interactions with the conserved residues in TM1 and TM7 including Y123^1.36b^, K127^1.40b^, Y130^1.43b^, I357^7.39b^ and L361^7.43b^. Consistently, the substitution of the residue at 1.36b by alanine greatly reduced PACAP27 potency for three receptors (136-fold for VIP2R, 15-fold for VIP1R, and 48-fold for PAC1R) (Fig. 3d). The VIP2R mutant I357^7.39b^A significantly diminished the PACAP27 potency by 69-fold (Fig. 3d), while equivalent mutations in VIP1R (M370^7.39b^A) and PAC1R (L382^7.39b^A) only displayed moderate effects (5-fold for VIP1R and 8-fold for PAC1R) (Supplementary Table 3). Besides the above common interactions among VIP2R, VIP1R and PAC1R, distinct amino acids in the equivalent positions of these three receptors fine tune the specific ligand-receptor recognition pattern. S2^P^ forms one hydrogen bond with Q356^7.38b^ of VIP2R, such an interaction is neither found in VIP1R (K369^7.38b^) nor in PAC1R (R381^7.38b^). By using leucine at position 3.36b instead of phenylalanine in both VIP1R and PAC1R, the contact between G4^P^ and VIP2R is slightly reduced. Consistently, the VIP2R mutant L209^3.36b^F increased the potency of PACAP27 by 3-fold (Fig. 3d). A similar phenomenon was observed for Y13^P^, where T136^1.33b^ in VIP1R and Y148^1.34b^ in PAC1R additionally provide one hydrogen bond and one stacking interaction, respectively. However, R12^P^ forms a salt bridge with D276 in the ECL2 of VIP2R, which is not observed in VIP1R, mainly due to the lack of negatively charged residues caused by a shorter ECL2 (by one amino acid compared to VIP2R or PAC1R).

The second network stabilizes the peptide–ECD–ECL1–TM1 interface through massive nonpolar and polar interactions. The C terminus of PACAP27 occupies a complementary binding groove of the ECD, consisting of a series of hydrophobic residues (I59^ECD^, F79^ECD^, F106^ECD^ and Y111^ECD^) that make extensive hydrophobic contacts with PACAP27 via V19^P^, F22^P^, L23^P^, V26^P^ and L27^P^, consistent with that seen in other class B1 GPCRs such as GLP-1R^24^, GHRHR^18^ and PAC1R^14^. Indeed, ECD deletion completely abolished the action of PACAP27, suggesting an essential role of the ECD (Fig. 3d). For the polar contacts, R14^P^ forms one hydrogen bond with the backbone atom of L183^2.71b^ and stacking interactions with Y184^ECL1^, the side-chain of Q16^P^ extends to the stalk with the formation of one hydrogen bond with N81^ECD^, while K20^P^ points to two adjacent negatively charged residues (D116^1.29b^ and E117^1.30b^) in the extracellular tip of TM1. These observations received support of our mutagenesis studies, where mutant F79^ECD^A, and Y184^ECL1^A decreased the potency of PACAP27-induced cAMP signaling by 76-fold, and 82-fold, respectively (Fig. 3d).

The most profound structural feature resides in the upper half of the PACAP27-bound VIP2R showing a position and orientation of the ECD N-terminal α-helix distinctive from all available class B1 GPCR structures reported to date (Figs. 3a, c and 4). Specifically, the tip of the N-terminal α-helix moved down toward the TMD by 9.8 Å relative to that of PAC1R (measured by the Cα of R26 in VIP2R and D23 in PAC1R) and inserted into a cleft between PACAP27 and the ECL1 (Fig. 4a). Such a unique conformation was probably caused by an outward movement of the ECL1, which appears to be conformationally more flexible as it is longer by three amino acids with the presence of two proline residues (P193^ECL1^ and P196^ECL1^) compared to the ECL1 of VIP1R or PAC1R. The inserted N-terminal α-helix stabilizes the peptide and ECL1 conformations via multiple contacts including one salt bridge (K15^P^ and E30^ECD^), one hydrogen bond (H28^ECD^ and T189^ECL1^) and several hydrophobic contacts (K15^P^ and F27^ECD^, Y22^P^ and I31 ^ECD^, L29^P^ and W199^ECL1^). Consistently, MD simulations found that the ECD intimately interacted with the peptide and ECL1 via the N-terminal α-helix (Supplementary Fig. 4). To reveal functional roles of the N-terminal α-helix in the presence of PACAP27, we truncated the N-terminal α-helix in a systemic manner and measured cAMP responses subsequently (Fig. 4b). For VIP2R, truncation of the ECD by five residues (VIP2R-Δ5) reduced PACAP27 potency by 2,874-fold, and cAMP signaling was completely abolished when ten or more residues were truncated (Fig. 4b and Supplementary Table 4). In contrast, the action of both VIP1R and PAC1R do not require the participation of the N-terminal α-helix or even the ECD, whose maximal responses of receptor-mediated cAMP accumulation in the presence of PACAP27 were retained when 5 or 10 residues even the entire ECD were truncated (Supplementary Table 4). A similar phenomenon was observed on dose-response characteristics of the N-terminal α-helix extension. Addition of flexible linker (G/S) at the receptor N terminus had neglectable effects on ligand binding and receptor activation for both VIP1R and PAC1R (Supplementary Table 4). However, VIP2R was extremely sensitive to the N-terminal α-helix extension, where two, five or ten introduced amino acids (G/S) reduced PACAP27 potency by 143-fold, 305-fold and 328-fold, respectively (Fig. 4b), indicating a curial role of the N-terminal α-helix length in VIP2R functioning. Collectively, our results suggest that VIP2R possesses a distinct molecular mechanism for peptide recognition and receptor activation.

**Figure 4.**
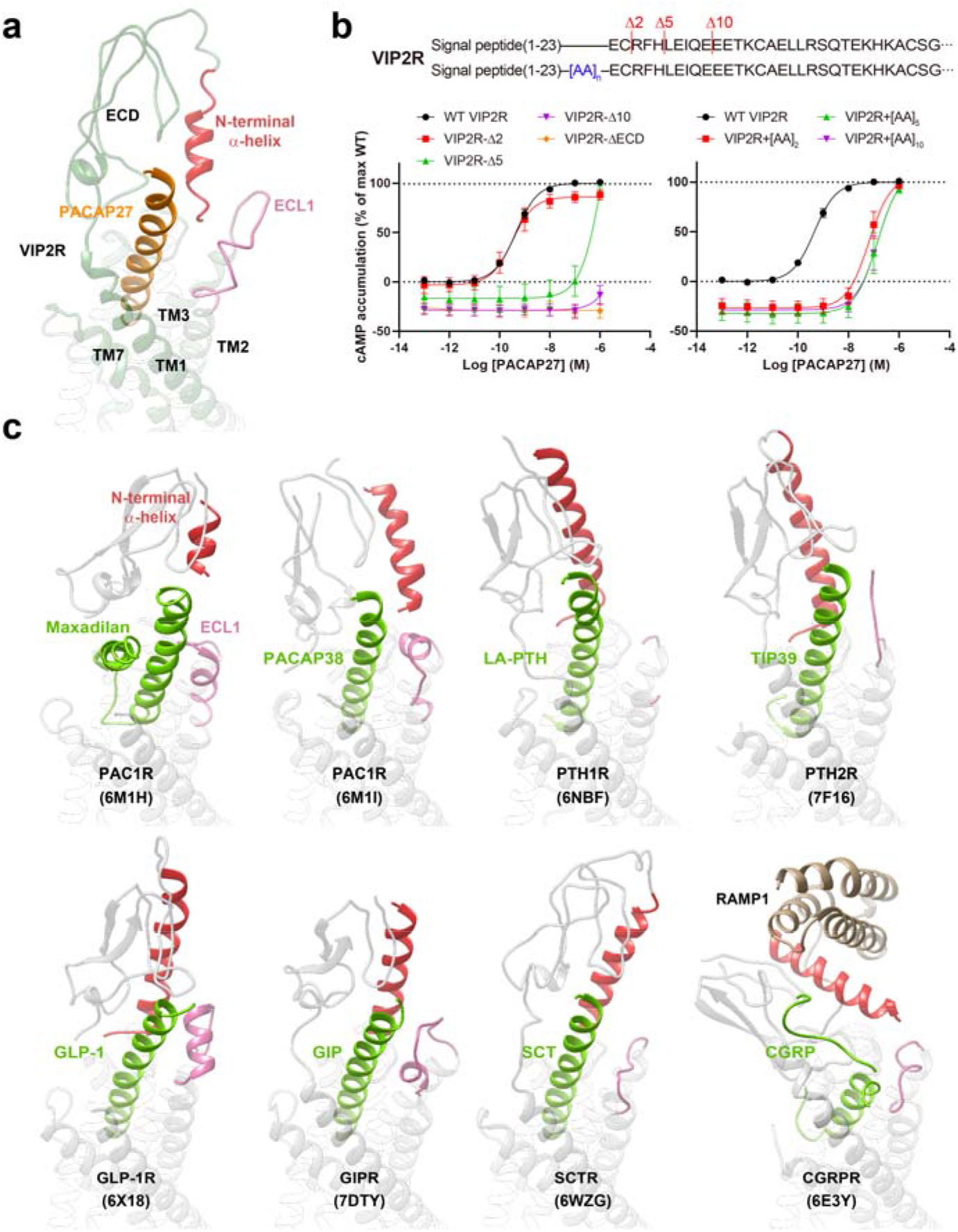
Unique conformation of the N-terminal α-helix of VIP2R in class B1 GPCRs. **a**, Close-up view of the PACAP27-ECD-ECL1 interface shows that the N-terminal α-helix filled the cleft between PACAP27 and ECL1, thereby stabilizing the complex. **b**, Signaling profiles of ECD-truncated or extended VIP2R in response to PACAP27. Data are presented as means ± S.E.M. of three independent experiments. WT, wild-type. Δ, residue truncation. [AA]_2_, extend the receptor N terminus by two amino acids (GS); [AA]_5_, five amino acids (GSSGG); [AA]_10_, ten amino acids (GSSGGGGSGG). **c**, Conformational comparison of the N-terminal α-helix among peptide-bound class B1 GPCR structures. All structures are superimposed on the GLP-1-bound GLP-1R (PDB code: 6×18)^24^ using the Cα carbons of the TMD residues. The receptor is shown in gray, peptide in green, ECL1 in pink and the N-terminal α-helix in red.

Inspired by such a unique N-terminal α-helix conformation of VIP2R, we performed structural analysis across class B1 GPCRs (Fig. 4c). Despite a high sequence similarity to VIP2R, PAC1R pulls out its N-terminal α-helix away from the peptide-ECL1 cleft to make negligible contact with either peptide or ECL1^14^. Alternatively, to hold PACAP38 or maxadilan (a native peptide from the sand fly), PAC1R adjusts its ECD and ECL1 conformation in a ligand-dependent manner, indicative of a high structural adaptability of PAC1R. In the case of parathyroid hormone (PTH) receptors whose ECL1s are unstructured, PTH1R^21^ and PTH2R^26^ rotate their N-terminal α-helices to stand upwards in line with the bound peptides, thereby providing additional contacts with the latter. Different from the ECL1-top conformation seen in PAC1R or the ECL2-top position observed in PTH1R/PTH2R, the N-terminal α-helices of glucagon receptor family members (GLP-1R^24^, GIPR^25^ and sCTR^28^ shown in Fig. 4c) locate in the middle of ECL1 and ECL2, and stabilize the peptide C terminus with the assistance of ECL1. As for calcitonin gene-related peptide receptor (CGRPR)^28^, the N-terminal α-helix rotates downward to cover the orthosteric site, probably due to the loop conformation in the C-terminal region of GCRP, as well as a shorter ECL1 compared to VIP2R or GLP-1R. Taken together, these observations demonstrate the diversity and flexibility of N-terminal α-helices among class B GPCRs, and highlight the importance of interplay among N-terminal α-helix, ECL1 and peptide.

### G protein coupling

Superimposing the TMD of G_s_-coupled VIP2R with that of VIP1R or PAC1R revealed a similar G protein-binding pocket created by the outward movement of the intracellular portion of TM6. Unsurprisingly, such movement is trigged by the formation of a ∼90° sharp kink at the middle of TM6 around the Pro^6.47b^-X-X-Gly^6.50b^ motif. Meanwhile, facilitated by G364^7.46b^ and G368^7.50b^ located in the middle of TM7, the extracellular half of TM7 bends towards TM6 to accommodate the entrance of the peptide N terminus. Besides these TM level conformational changes, that of the residue level including the rearrangement of the central polar network, HETX motif and TM2-6-7-helix 8 polar network were also seen in VIP2R, in line with previous observations among class B1 GPCRs^14, 18, 24, 29^.

As shown in Fig. 5, G_s_ protein is anchored by the α5 helix of Gα_s_ (GαH5), thereby fitting to the cytoplasmic cavity formed by TMs 2, 3, 5, 6, 7 and ICLs 1–3, while Gβ mainly interacts with H8 of VIP2R (Fig. 5b-c). Specially, R325^6.37b^, S329^6.41b^ and S380^8.48b^ form multiple hydrogen bonds with the C terminus of GαH5, namely, L394^GαH5^ (via carboxy terminus), L393^GαH5^ (via backbone atom) and E392^GαH5^(via side-chain atom), respectively. In addition, Y391^GαH5^, L393^GαH5^and L394^GαH5^ contribute massive hydrophobic contacts with the hydrophobic residues in TMs 3, 5 and 6, such as L227^3.54b^, L231^3.58b^, I302^5.57b^, L306^5.61b^, K309^5.64b^, L332^6.44b^ and L333^6.45b^. There are some receptor-specific structural features displayed by the ICL2. Different from the benzene ring of F248^ICL2^ (VIP1R) and F259^ICL2^ (PAC1R) that inserted into a hydrophobic pocket of GαH5, M234^ICL2^ of VIP2R (M234^ICL2^) makes slightly reduced interactions with Gα_s_ (Fig. 5d). However, the dipped down side-chain conformation of L235^ICL2^ provides additional hydrophobic contacts to stabilize the VIP2R–G_s_ interface, while that of VIP1R (F249^ICL2^) or PAC1R (F260^ICL2^) rotates away from the interface that has negligible contact with G protein (Fig. 5e).

**Figure 5.**
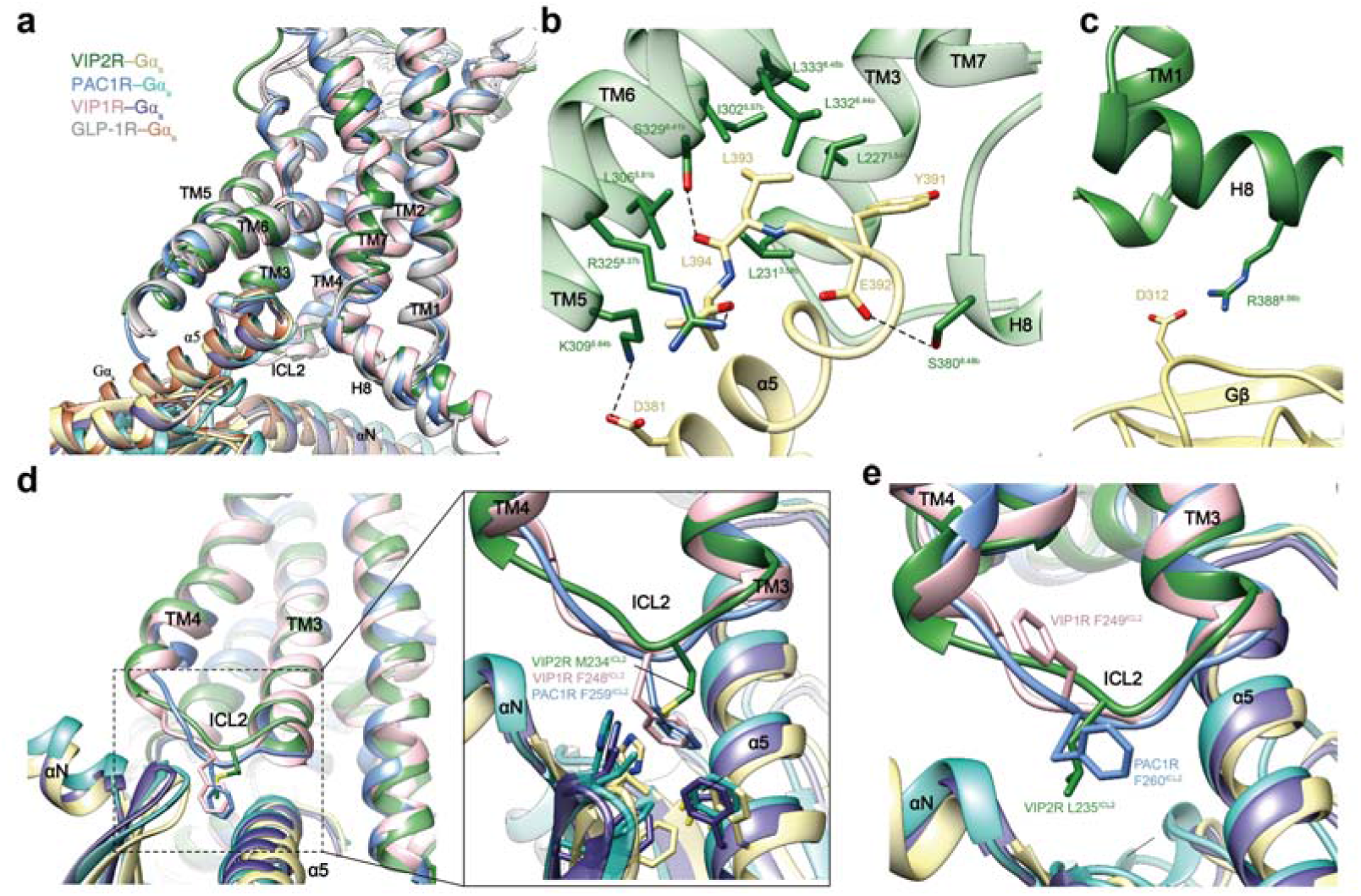
G protein coupling of VIP2R. **a**, Comparison of G protein coupling among VIP2R, VIP1R (PDB code: 6VN7), PAC1R (PDB code: 6P9Y)^16^ and GLP-1R (PDB code: 6×18)^24^. The receptors and G proteins are colored as the labels. **b**, Interaction between VIP2R and the C terminus of Gα_s_. **c**, Interactions between H8 and Gβ of VIP2R. **d-e**, Comparison of the interactions between ICL2 and Gα_s_ for VIP2R, VIP1R and PAC1R. Residues involved in the interactions are shown as sticks.

## Discussion

The cryo-EM structure of the PACAP27–VIP2R–G_s_ complex presented here reveals a distinctive and previously unknown peptide recognition mechanism responsible for ligand specificity at near-atomic resolution. Combined with functional and MD simulation studies, we determined that PACAP27 is recognized by VIP2R through its N-terminal α-helix that inserts into the cleft between the peptide and the ECL1. Such a phenomena was only observed in VIP2R, not in closely related VIP1R and PAC1R, indicating a diversified receptor responsiveness to the same ligand. The importance of receptor N terminus was also elegantly demonstrated in a recent study on compound 2-bound GLP-1R showing another TMD-interacted conformation for the N-terminal α-helix, which penetrates to the GLP-1 binding site and activate the receptor^19^. The interplay between N-terminal α-helix and TMD observed in different class B1 receptors support the notion that the N-terminal α-helix (broadly the ECD region) is involved in regulating receptor-mediated signal transduction in addition to its constitutive role of binding to the peptide C terminus. Obviously, the physiological significance of these observations require in-depth investigations including the role of ECD in basal activities of certain class B1 receptors. Together with 14 other class B1 GPCRs having full-length structures, addition of the PACAP27-bound VIP2R–G_s_ complex structure will allow us to perform a class-wide analysis and comparison of their structural and functional features with a goal of developing better therapeutic agents for a variety of human diseases.

## Methods

### Construct

Human VIP2R (residues 24-438) was cloned into pFastBac (Invitrogen) with an N-terminal FLAG tag followed by a His8 tag. A TEV protease cleavage site followed by a double maltose binding protein (2MBP) tag and LgBiT at the C terminus via homologous recombination (CloneExpress One Step Cloning Kit, Vazyme). The native signal peptide was replaced with the prolactin precursor sequence to increase the protein expression. A dominant negative bovine Gα_s_ (DNGα_s_) construct was generated by site-directed mutagenesis to incorporate mutations S54N, G226A, E268A, N271K, K274D, R280K, T284D, I285T and A366S to decrease the affinity of nucleotide-binding and increase the stability of Gαβγ complex^30^. Rat Gβ1 was cloned with an N-terminal His6 tag and a C-terminal SmBiT connected with a 15-residue linker. All three G protein components together with bovine Gγ2 were cloned into a pFastBac vector, respectively.

### Cell culture

Spodoptera frugiperda (*Sf*9) insect cells (Expression Systems) were cultured in ESF 921 serum-free medium (Expression Systems) at 27°C and 120 rpm. Cell cultures were grown to a density of 2.5 × 10^6^ cells mL^−1^ and then infected with baculoviruses expressing VIP2R–LgBiT fusion, DNGα_s_, Gβ1–SmBiT fusion and Gγ2, respectively, at the ratio of 1:2:2:2. The cells were collected by centrifugation at 2000 rpm for 20 min after infection for 48 h, and kept frozen at -80°C until use.

### PACAP27–VIP2R–G_s_ complex formation and purification

Cell pellets from 1 L culture were thawed and lysed in the lysis buffer (20 mM HEPES, pH 7.4, 100 mM NaCl, 10% (v/v)). The complex formation was initiated by addition of 10 μM PACAP27 (Synpeptide), 20 μg/mL Nb35, 25 mU/mL apyrase (NEB), 5 mM CaCl_2_, 5 mM MgCl_2_ and 250 μM TCEP, supplemented with EDTA-free protease inhibitor cocktail (Bimake) for 1.5 h incubation at room temperature (RT). The membrane was solubilized by 0.5% (w/v) lauryl maltose neopentyl glycol (LMNG; Anatrace) and 0.1% (w/v) cholesterol hemisuccinate (CHS; Anatrace) for 2 h at 4°C. After centrifugation at 65,000× *g* for 40 min, the supernatant was separated and incubated with amylose resin (NEB) for 2 h at 4°C. The resin was collected and packed into a gravity flow column and washed with 20 column volumes of 5 μM PACAP27, 0.01% (w/v) LMNG, 0.002% (w/v) CHS, 0.01% (w/v) GDN, 0.008% (w/v) CHS, 20 mM HEPES, pH7.4, 100 mM NaCl, 10% (v/v) glycerol, 2 mM MgCl_2_, 2 mM CaCl_2_ and 25 μM TCEP. 2MBP-tag was removed by His-tagged TEV protease (customer-made) during overnight incubation. The complex was concentrated using an Amicon Ultra Centrifugal filter (MWCO, 100 kDa) and subjected to a Superose 6 Increase 10/300 GL column (GE Healthcare) that was pre-equilibrated with running buffer containing 20 mM HEPES, pH 7.4, 100 mM NaCl, 2 mM MgCl_2_, 2 mM CaCl_2_, 250 μM TCEP, 5 μM PACAP27, 0.00075% (w/v) LMNG, 0.00025% (w/v) GDN, 0.00025% digitonin and 0.0002% (w/v) CHS. Eluted fractions containing the PACAP27**–**VIP2R**–**G_s_ complex were pooled and concentrated. All procedures mentioned above were performed at 4°C.

### Expression and purification of Nb35

The nanobody 35 (Nb35) with a C-terminal histidine tag (His6) was expressed in *E. coli* BL21 (DE3) bacteria and cultured in TB medium supplemented with 2 mM MgCl_2_, 0.1% (w/v) glucose and 50 μg/mL ampicillin to an OD600 value of 0.7-1.2 at 37°C. The culture was then induced by 1 mM IPTG and grown overnight incubation at 28°C. Cells were harvested by centrifugation (4000 rpm, 20 min) and Nb35 protein was extracted and purified by nickel affinity chromatography as previously described^31^. Eluted protein was concentrated and subjected to a HiLoad 16/600 Superdex 75 column (GE Healthcare) pre-equilibrated with buffer containing 20 mM HEPES, pH 7.5 and 100 mM NaCl. The monomeric fractions supplemented with 30% (v/v) glycerol were flash frozen in liquid nitrogen and stored in -80°C until use.

### Cryo-EM data acquisition

The purified PACAP27**–**VIP2R**–**G_s_ complex (3 μL at 3.7 mg per mL) was applied to glow-discharged holey carbon grids (Quantifoil R1.2/1.3). Vitrification was performed using a Vitrobot Mark IV (ThermoFisher Scientific) at 100% humidity and 4°C. Cryo-EM images were processed on a Titan Krios microscope (FEI) equipped with a Gatan K3 Summit direct electron detector. The microscope was operated at 300 kV accelerating voltage, at a nominal magnification of 46,685× in counting mode, corresponding to a pixel size of 1.071Å. In total, 4,753 movies were obtained with a defocus range of -1.2 to -2.2 μm. An accumulated dose of 80 electrons per Å^2^ was fractionated into a movie stack of 36 frames.

Dose-fractionated image stacks were subjected to beam-induced motion correction using MotionCor2.1. A sum of all frames, filtered according to the exposure dose, in each image stack was used for further processing. Contrast transfer function parameters for each micrograph were determined by Gctf v1.06. Particle selection, 2D and 3D classifications were performed on a binned dataset with a pixel size of 2.142 Å using RELION-3.1.1. Auto-picking yielded 5,558,869 particle projections that were subjected to reference-free 2D classification to discard false positive particles or particles categorized in poorly defined classes, producing 1,729,374 particle projections for further processing. This subset of particle projections was subjected to a round of maximum-likelihood-based 3D classifications with a pixel size of 2.142 Å, resulting in one well-defined subset with 931,248 projections. Further 3D classifications with mask on the receptor produced one good subset accounting for 602,466 particles, which were subjected to another round of 3D classifications with mask on the ECD. A selected subset containing 305,004 projections was then subjected to 3D refinement and Bayesian polishing with a pixel size of 1.071 Å. After the last round of refinement, the final map has an indicated global resolution of 3.4 Å at a Fourier shell correlation (FSC) of 0.143. Local resolution was determined using the Bsoft package with half maps as input maps.

### Model building and refinement

The model of the PACAP27–VIP2R–Gs complex was built using the cryo-EM structure of PACAP27–VIP1R–G_s_ complex (PDB code: 6VN7) and the crystal structure of VIP2R ECD (PDB code: 2×57) as the starting point. The model was docked into the EM density map using Chimera^32^, followed by iterative manual adjustment and rebuilding in COOT^33^. Real space refinement was performed using Phenix^34^. The model statistics were validated using MolProbity^35^. Structural figures were prepared in Chimera and PyMOL (https://pymol.org/2/). The final refinement statistics are provided in Supplementary Table 1.

### Molecular dynamics simulations

Molecular dynamic simulations were performed by Gromacs 2020.1. The peptide–VIP2R complexes were built based on the cryo-EM structure of the PACAP27–VIP2R–G_s_ complex and prepared by the Protein Preparation Wizard (Schrodinger 2017-4) with the G protein and Nb35 nanobody removed. The receptor chain termini were capped with acetyl and methylamide. All titratable residues were left in their dominant state at pH 7.0. To build MD simulation systems, the complexes were embedded in a bilayer composed of 254 POPC lipids and solvated with 0.15 M NaCl in explicit TIP3P waters using CHARMM-GUI Membrane Builder v3.5^36^. The CHARMM36-CAMP force filed^37^ was adopted for protein, peptides, lipids and salt ions. The Particle Mesh Ewald (PME) method was used to treat all electrostatic interactions beyond a cut-off of 10Å and the bonds involving hydrogen atoms were constrained using LINCS algorithm^38^. The complex system was first relaxed using the steepest descent energy minimization, followed by slow heating of the system to 310 K with restraints. The restraints were reduced gradually over 50 ns. Finally, restrain-free production run was carried out for each simulation, with a time step of 2 fs in the NPT ensemble at 310 K and 1 bar using the Nose-Hoover thermostat and the semi-isotropic Parrinello-Rahman barostat^39^, respectively.

### cAMP accumulation assay

Wild-type (WT) or mutant VIP2Rs, VIP1Rs and PAC1Rs were cloned into pcDNA3.1 vector (Invitrogen) for functional studies. CHO-K1 cells were transiently transfected with the vectors using Lipofectamine 2000 transfection reagent (Invitrogen) and incubated at 37°C in 5% CO_2_. After 24 h, the transfected cells were digested with 0.02% (w/v) EDTA, resuspended in stimulation buffer (Hanks’ balanced salt solution (HBSS) supplemented with 5 mM HEPES, 0.5 mM IBMX and 0.1% (w/v) BSA, pH 7.4) to a density of 0.6 million cells per mL and added to 384-well white plates (3000 cells per well). cAMP accumulation was measured by a LANCE Ultra cAMP kit (PerkinElmer) according to the manufacturer’s instructions. In brief, transfected cells were incubated for 40 min in stimulation buffer with different concentrations of ligand (5 μL) at RT. The reaction was stopped by addition of lysis buffer containing 5 μL Eu-cAMP tracer and 5 μL ULight-anti-cAMP. Plates were then incubated for 60 min at RT and time-resolved FRET signals were measured at 620 nm and 665 nm, respectively, by an EnVision multilabel plate reader (PerkinElmer). Data were analyzed in GraphPad PRISM 8 and all values were normalized to the WT for each ligand.

### Whole cell binding assay

CHO-K1 cells were cultured in F12 medium with 10% FBS and seeded at a density of 30,000 cells/well in Isoplate-96 plates (PerkinElmer). Twenty-four hours after transfection with the WT or mutant receptors, CHO-K1 cells were washed twice and incubated with blocking buffer (F12 supplemented with 25 mM HEPES and 0.1% (w/v) BSA, pH 7.4) for 2 h at 37°C. For homogeneous competition binding, radiolabeled ^125^I-PACAP27 (40 pM, PerkinElmer) and seven decreasing concentrations of unlabeled peptides were added separately and competitively reacted with the cells in blocking buffer at RT for 3 h. Following incubation, cells were washed three times with ice-cold PBS and lysed by 50 μL lysis buffer (PBS supplemented with 20 mM Tris-HCl, 1% Triton X-100, pH 7.4). The radioactivity was subsequently counted (counts per minute, CPM) in a scintillation counter (MicroBeta2 Plate Counter, PerkinElmer) using a scintillation cocktail (OptiPhase SuperMix, PerkinElmer).

## Data availability

All relevant data are available from the corresponding authors upon reasonable request. The raw data underlying Figs. 1b, 3d, 4b, Supplementary Figs. 1c-d, and Supplementary Tables 3-4 are provided as a Source Data file. The atomic coordinate and electron microscopy map of the PACAP27–VIP2R–G_s_ complex have been deposited in the Protein Data Bank (PDB) under accession code: XXX and Electron Microscopy Data Bank (EMDB) accession code: EMD-XXX, respectively. Source data are provided with this paper.

## Acknowledgements

We thank E. Zhou and F.L. Zhou for technical advice. This work was partially supported by National Natural Science Foundation of China 81872915 (M.-W.W.), 82073904 (M.-W.W.), 81773792 (D.Y.), 81973373 (D.Y.) and 21704064 (Q.T.Z.); National Science & Technology Major Project of China–Key New Drug Creation and Manufacturing Program 2018ZX09735–001 (M.-W.W.) and 2018ZX09711002–002–005 (D.Y.); the National Key Basic Research Program of China 2018YFA0507000 (M.-W.W.), Shanghai Municipal Science and Technology Major Project 2019SHZDZX02 (H.E.X.); Ministry of Science and Technology of China Major Project XDB08020303 (H.E.X.); Novo Nordisk-CAS Research Fund grant NNCAS-2017–1-CC (D.Y.); The Young Innovator Association of CAS Enrollment (L.H.Z.) and SA-SIBS Scholarship Program (L.H.Z. and D.Y.). The cryo-EM data were collected at Cryo-Electron Microscopy Research Center, Shanghai Institute of Materia Medica.

## Author contributions

Y.N.X. designed the expression constructs, purified the receptor complexes, screened specimen, prepared the final samples for negative staining/data collection towards the structure and participated in manuscript preparation with the help of W.B.F., F.H.Z. and Y.J.; Y.N.X., W.B.F. and J.L. performed functional studies; A.T.D. conducted ligand binding assay; A.Y.L. T.X. and Q.T.Z. performed map calculation, model building and figure preparation; Q.T.Z. carried out MD simulations; Q.T.Z. and L.H.Z. conducted structural analysis; D.Y. oversaw mutagenesis and signaling experiments, participated in data analysis and manuscript preparation; H.E.X. and M.-W.W. initiated the project, supervised the studies and analyzed the data; Q.T.Z. and M.-W.W. wrote the manuscript with inputs from all co-authors.

## Competing interests

The authors declare no competing interests.

## Additional information

**Supplementary information** is available for this paper at XXXX.

Correspondence and requests for materials should be addressed to H.E.X., D.Y. or M.-W.W.

---

**Supplementary Figure 1.**
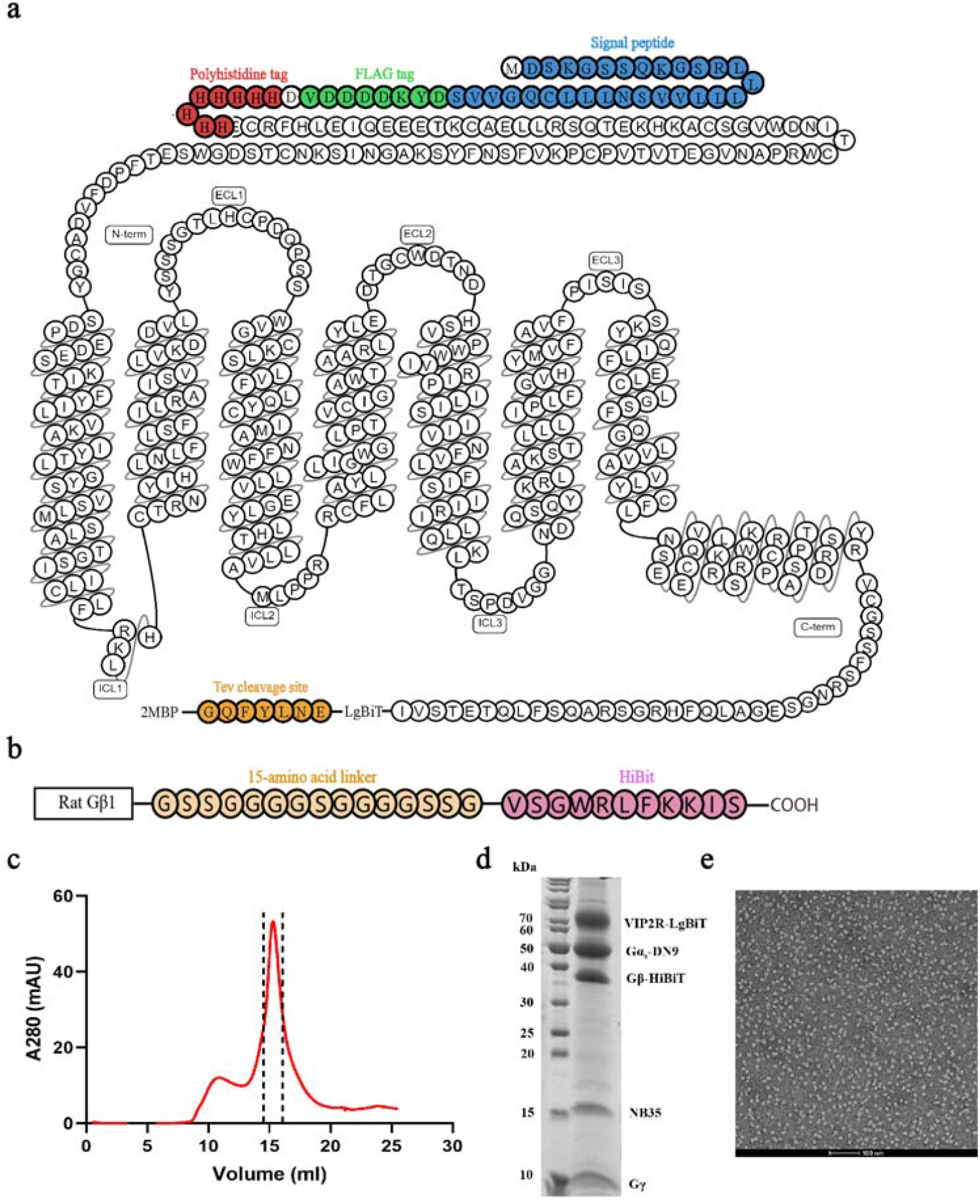
Purification and characterization of the PACAP27–VIP2R–G_s_–Nb35 complex. **a**, Snake-plot diagram of the human VIP2R**–**LgBiT construct. **b**, Gβ1 constructs used for structure determination. Rat Gβ1 was attached to HiBiT with a 15-amino acid (15AA) linker between them. **c-e**, Analytical size-exclusion chromatography (**c**), SDS-PAGE/Coomassie blue stain (**d**) and representative negative staining image (**e**) of the purified PACAP27–VIP2R–G_s_ complex.

**Supplementary Figure 2.**
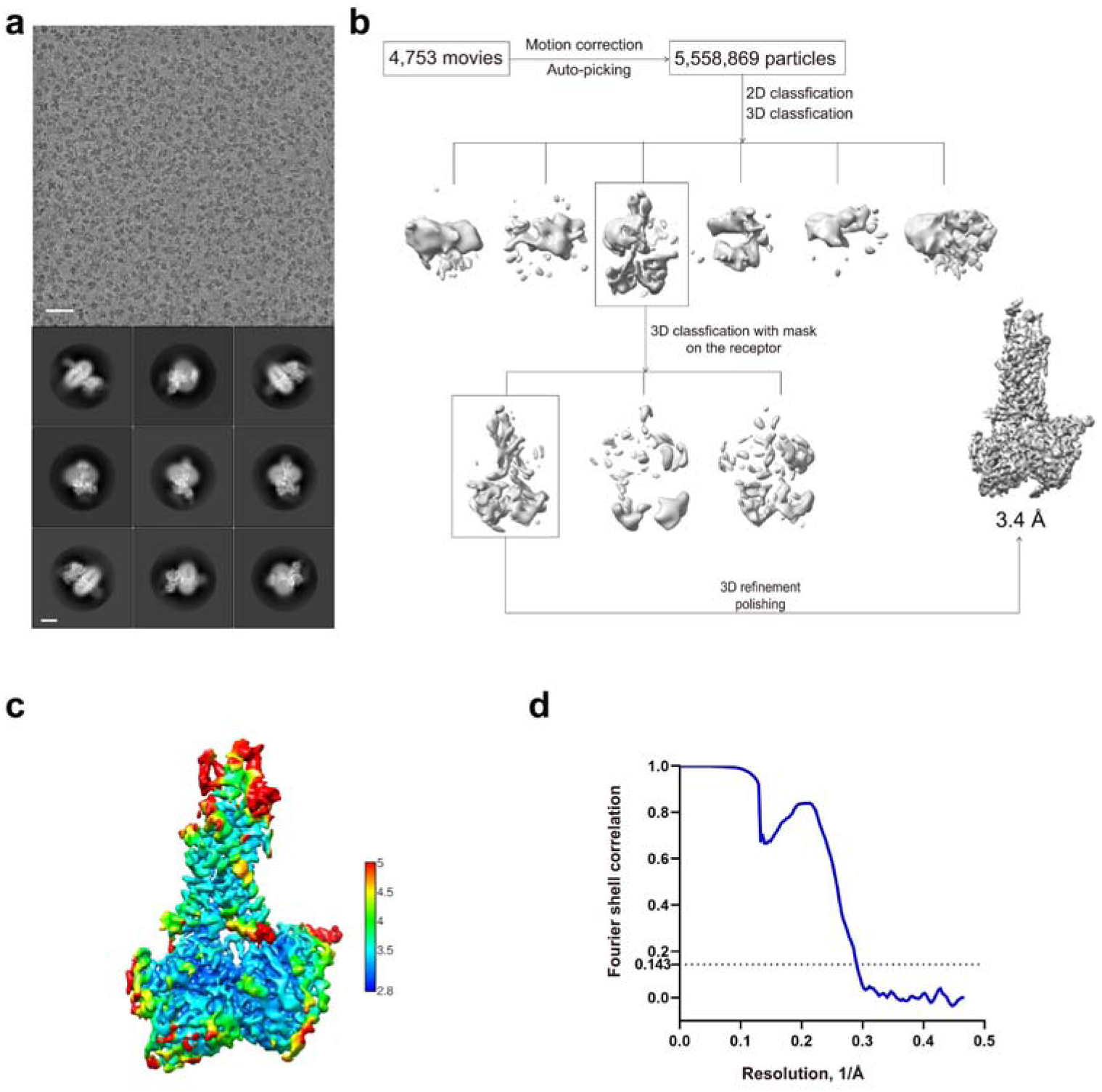
Cryo-EM analysis of the PACAP27–VIP2R–G_s_ complex. **a**, representative cryo-EM micrograph (scale bar: 40 nm) and two dimensional class averages (scale bar: 5 nm). **b**, flowchart of cryo-EM data processing. **c**, local resolution distribution map of the PACAP27–VIP2R–G_s_ complex. **d**, Gold-standard Fourier shell correlation (FSC) curve of overall refined model.

**Supplementary Figure 3.**
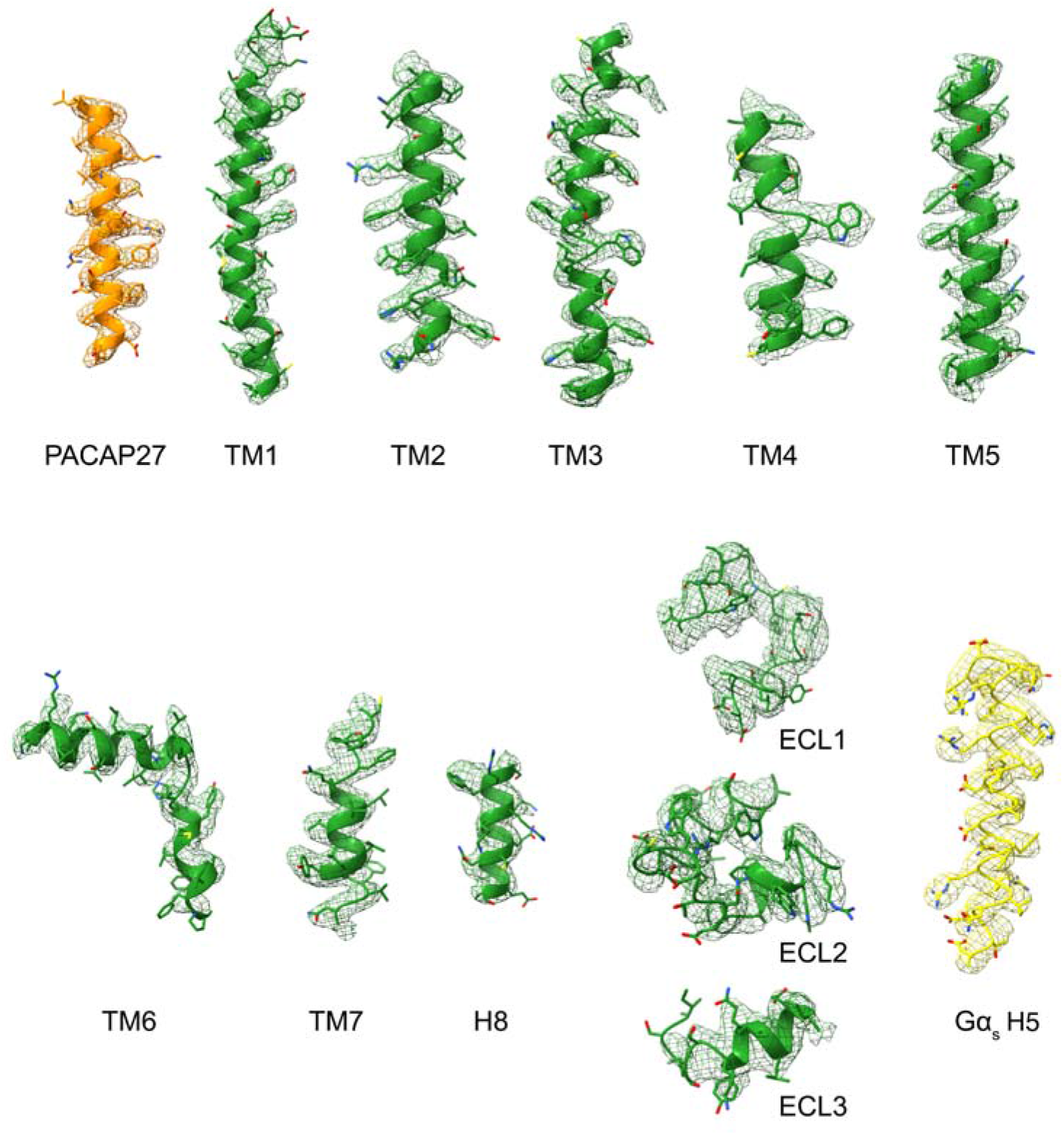
Atomic resolution model of the PACAP27–VIP2R–G_s_ complex in the cryo-EM density map. EM density map and model are shown for all seven transmembrane α-helices, helix 8 and all extracellular loops of VIP2R, the α5-helix (H5) of the Gα_s_ Ras-like domain and PACAP27.

**Supplementary Fig 4.**
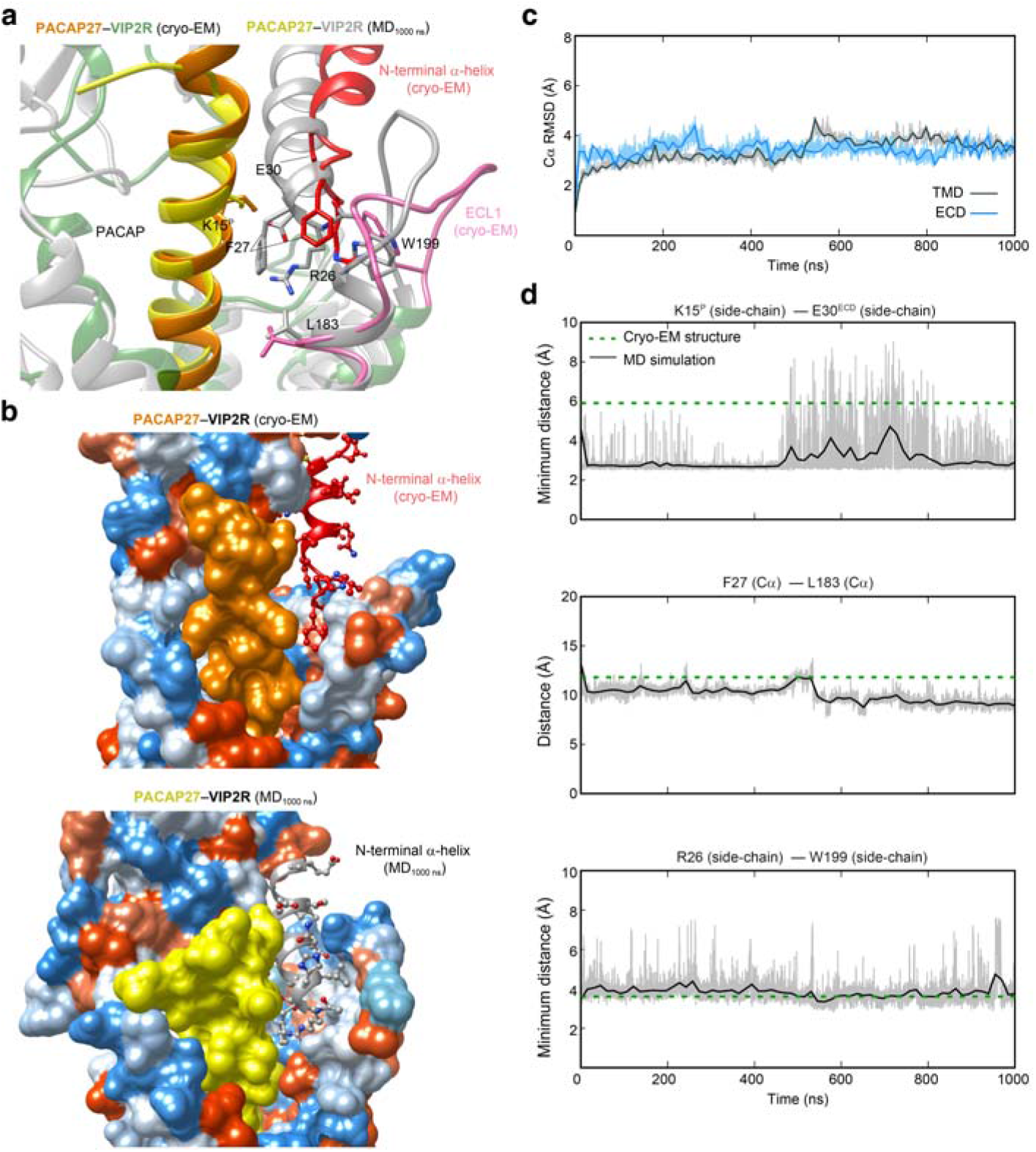
Molecular dynamics (MD) simulations of PACAP27-bound active VIP2R. **a**, Comparison of the N-terminal α-helix conformation between simulation snapshot and the cryo-EM structure. The key residues in the peptide-receptor interface are shown in sticks. **b**, Surface representation of the peptide-ECL1 cleft that N-terminal α-helix inserted for the cryo-EM structure (top panel) and finial MD snapshot at 1000 ns (bottom panel). The receptor is shown in surface representation and colored from dodger blue for the most hydrophilic region, to white, to orange red for the most hydrophobic region. **c**, Root mean square deviation (RMSD) of Cα positions of the VIP2R ECD and TMD, where all snapshots were superimposed on the cryo-EM structure of VIP2R ECD and TMD using the Cα atoms, respectively. **d**, Representative minimum distances between the N-terminal α-helix and peptide or TMD (Top, K15^P^ - E30 ; middle, F27^ECD^ - L183^2.71b^; bottom, R26^ECD^ - W199^ECL1^).

## Notes

### Competing Interest Statement

The authors have declared no competing interest.

